# Overexpression of the chloroplastic 2-oxoglutarate/malate transporter in rice disturbs carbon and nitrogen homeostasis

**DOI:** 10.1101/2020.03.31.018226

**Authors:** Shirin Zamani-Nour, Hsiang-Chun Lin, Berkley J. Walker, Tabea Mettler-Altmann, Roxana Khoshravesh, Shanta Karki, Efren Bagunu, Tammy L. Sage, W. Paul Quick, Andreas P.M. Weber

**Author notes:** These authors contributed equality to this paper (Shirin Zamani-Nour, Hsiang-Chun Lin). Correspondence Andreas P.M. Weber Institute of Plant Biochemistry, Heinrich-Heine-University, Universitätsstraße 1, D-40225 Düsseldorf.

## Abstract

The chloroplastic oxaloacetate/malate transporter (OMT1 or DiT1) takes part in the malate valve that protects chloroplasts from excessive redox poise through export of malate and import of oxaloacetate (OAA). Together with the glutamate/malate transporter (DCT1 or DiT2), it connects carbon with nitrogen assimilation, by providing α-ketoglutarate for the GS/GOGAT reaction and exporting glutamate to the cytoplasm. OMT1 further plays a prominent role in C_4_ photosynthesis. OAA resulting from PEP-carboxylation is imported into the chloroplast, reduced to malate by plastidic NADP-MDH, and then exported for transport to bundle sheath cells. Both transport steps are catalyzed by OMT1, at the rate of net carbon assimilation. Therefore, to engineer C_4_ photosynthesis into C_3_ crops, OMT1 must be expressed in high amounts on top of core C_4_ metabolic enzymes. We report here high-level expression of *ZmOMT1* from maize in rice (*Oryza sativa* ssp. *indica* IR64). Increased activity of the transporter in transgenic rice was confirmed by reconstitution of transporter activity into proteoliposomes. Unexpectedly, over-expression of *ZmOMT1* in rice negatively affected growth, CO_2_ assimilation rate, total free amino acid contents, TCA cycle metabolites, as well as sucrose and starch contents. Accumulation of high amounts of aspartate and the impaired growth phenotype of OMT1 rice lines could be suppressed by simultaneous over-expression of *ZmDiT2*. Implications for engineering C_4_-rice are discussed.

## Introduction

Population growth, climate change, and lack of arable land are placing greater dependence on crop yield improvement. However, crop demand is already outpacing the yield gains achieved by conventional breeding and hence, step-wise changes in crop yield are needed (Kromdijk and Long, 2016). Rice (*Oryza sativa* L.) is a C_3_ plant that belongs to the Gramineae family and is one of the most important staple crops in the world. Its highest consumption is in Asia (Muthayya *et al*., 2014) where 60% of the world population exists (Bai *et al*., 2018), with the highest and lowest rates of poverty and income, respectively (FAO, 2017). Therefore, boosting rice yield and performance is an important goal for improving the quality of life for a large share of the global population. Engineering the C_3_ crop rice to perform C_4_ photosynthesis would greatly improve rice productivity by up to 50% per year (Wang *et al*., 2016), through maximizing the conversion of the captured solar energy into chemical energy and biomass (Hibberd *et al*., 2008).

C_3_ photosynthesis performs both initial carbon fixation and Calvin-Benson cycle reactions in the mesophyll (M). In C_4_ photosynthesis, initial carbon fixation and the Calvin-Benson cycle are carried out separately in M and one or more layers of sheath cells (bundle and/or mestome sheath) surrounding the vascular tissue, respectively. This spatial separation concentrates CO_2_ around the enzyme Ribulose-1,5-bisphosphate carboxylase/oxygenase (RubisCO), thereby, reducing RubisCO oxygenase activity and the subsequent loss of energy and previously fixed CO_2_ during photorespiration (Sage *et al*., 2012). Suppressing the energy- and CO_2_-loss from photorespiration leads to greater plant biomass, nitrogen- and water-use efficiency (Ghannoum *et al*., 2011). C_4_ photosynthesis also represents an adaptation for coping with stressful conditions, such as drought, high temperature, and light intensity (Edwards *et al*., 2010).

Chloroplasts with their double-envelope membrane and internal compartments play a critical role in carbon fixation and photosynthesis. Since biological membranes form barriers for the diffusion of hydrophilic metabolites, membrane transporters are required for the selective flux of polar molecules and metabolites across the chloroplast membrane (Haferkamp and Linka, 2012). One of the transporters that resides in the plastid inner envelope membrane is known as <u>o</u>xaloacetate/<u>m</u>alate transporter <u>1</u> (OMT1) or <u>di</u>carboxylate transporter <u>1</u> (DiT1). The gene is expressed ubiquitously in roots, stems, leaves, florescence, and siliques of mature Arabidopsis plants (Taniguchi *et al*., 2002) and the protein is an oxaloacetate/malate antiporter with 12 α-helical transmembrane domains. OMT1/DIT1 functions to transport substrates according to the electrochemical gradient generated by solutes in and outside the chloroplast membrane (Weber *et al*., 1995). This transporter, in concert with malate dehydrogenase (plastidic and cytosolic isoforms), forms the malate shunt that plays a key role in exporting excess reducing compounds from the chloroplast, to protect photosystem II and to balance stromal redox potential (Selinski and Scheibe, 2019).

Redox balancing through the OMT1/DIT1-mediated malate valve is expected to be more beneficial when photorespiratory rates are increased, since higher relative rates of photorespiration increase the ATP/NAPDH demand of central metabolism resulting in an excess of reduced NADPH in the plastid (Kramer and Evans, 2011; Walker *et al*., 2014). The malate valve can serve to oxidize the over-reduced NADPH pool to regenerate oxidized NADP^+^ carriers that are needed to maintain electron transport. The NADH generated in the cytosol from the malate valve activity is consumed in other reactions such as nitrate reduction. The resulting nitrite is imported into chloroplasts where it is further reduced to ammonia that is subsequently assimilated into glutamate by the GS/GOGAT (glutamine synthase/glutamate synthase) pathway (Tobin and Yamaya, 2001, Selinski and Scheibe, 2019). Glutamate itself is a building block for the biosynthesis of many amino acids (Forde and Lea, 2007). OMT1, jointly with the DiT2/DCT1 transporter (a glutamate/malate transporter), therefore, connects carbon and nitrogen metabolism while equilibrating the ATP/NADPH ratio in chloroplast stroma (Taniguchi *et al*., 2002, Kinoshita *et al*., 2011, Taniguchi and Miyake, 2012). The strong, visibly perturbed phenotypes of *omt1* mutants in Arabidopsis (Kinoshita *et al*., 2011) and in tobacco (Schneidereit *et al*., 2006) confirm its crucial role in carbon and nitrogen assimilation pathways as well in plant growth and development.

OMT1 also plays an important role in C_4_ photosynthesis. The transporter imports oxaloacetate (OAA) that is formed by cytosolic phosphoenolpyruvate carboxylase (PEPC) into M cell chloroplasts where it is reduced to malate by NADP-malate dehydrogenase (NADP-MDH). OMT1 also facilitates the export of malate to the cytosol. These transport steps occur at the same rate as CO_2_ assimilation and thus, for engineering C_4_ photosynthesis into C_3_ crops such as rice, high expression and activity of OMT1 are required. In this study, as part of the effort to engineer C_4_ rice, we introduced the *ZmOMT1* gene from C_4_-maize into C_3_-rice to achieve sufficiently highly transport capacity for OAA and malate across the chloroplast envelope. Additionally, since C_4_ photosynthesis requires a complex array of biochemical and anatomical components, we investigate whether *ZmOMT1* expression triggers anatomical features found in C_4_ plants, which could further aid continued C_4_ engineering efforts.

## Materials and methods

### Rice transformation and growth conditions

To express maize *ZmOMT1* in rice M cells, we transferred the pSC110: *ZmOMT1*:AcV5 construct into *Oryza sativa* spp. *indica* cultivar IR64 (Mackill and Khush, 2018). The construct contains the full-length cDNA of the *ZmOMT1* gene (GRMZM2G383088) from maize (*Zea mays* var. B73) with a C-terminal AcV5 epitope tag, driven by the maize M-specific *ZmPEPC* promoter from pSC110 vector (Supplementary Fig. 1A). The Fwd_primer 5’-CACCATGGTCGACGCGTCCTCCAC and the Rev_primer 5’-TCAAGACCAGCCGCTCGCATCTTTCCAAGACCACAGCCCGATTATCTTC were used to clone the coding sequence via the pENTR vector into the pSC110 expression vector utilizing the Gateway cloning system (Thermo Fisher Scientific). The pSC110 vector was generated as previously described (Osborn *et al*., 2017). The AcV5 epitope tag was placed downstream of *ZmOMT1* coding sequence for later detection of expressed protein using commercially available AcV5 antibody. The final construct was transferred into freshly harvested immature embryos 8-12 days after anthesis using an *Agrobacterium*-mediated transformation protocol as described in Yin *et al*., (2019). After one week of co-cultivation and following 5 days on non-selective medium, emerging resistant calli were selected with 30 mg/l hygromycin B. The transgenic plants generated from hygromycin-resistant calli were transferred to Yoshida hydroponic solution (Yoshida *et al*., 1972) for 2 weeks and then transplanted into 0.5 L pots filled with soil. Plants were grown in a greenhouse at the International Rice Research Institute (IRRI, Los Baños, Philippines: 14°9’53.58’’S 121°15’32.19’’E). The average day/night temperatures were 35±3°C and 28±3°C, respectively. The average and maximum light intensities were 825 µmol photons m^−2^ s^−1^ and 2000 µmol photons m^−2^ s^−1^, respectively. Seeds of transgenic plants were germinated in distilled water for one week and transplanted into soil in 100 ml Rootrainers (http://rootrainers.co.uk/). After 2 weeks, plants were transplanted to 7 L soil pots. Plants were grown at Heinrich-Heine University (HHU) Düsseldorf, Germany under semi-controlled greenhouse conditions (16h day/8h night and 25°C). Assessment of leaf gas exchange, as well as metabolite, C:N ratio, total free amino acids, and transporter activity measurements were performed at HHU.

**Fig. 1:**
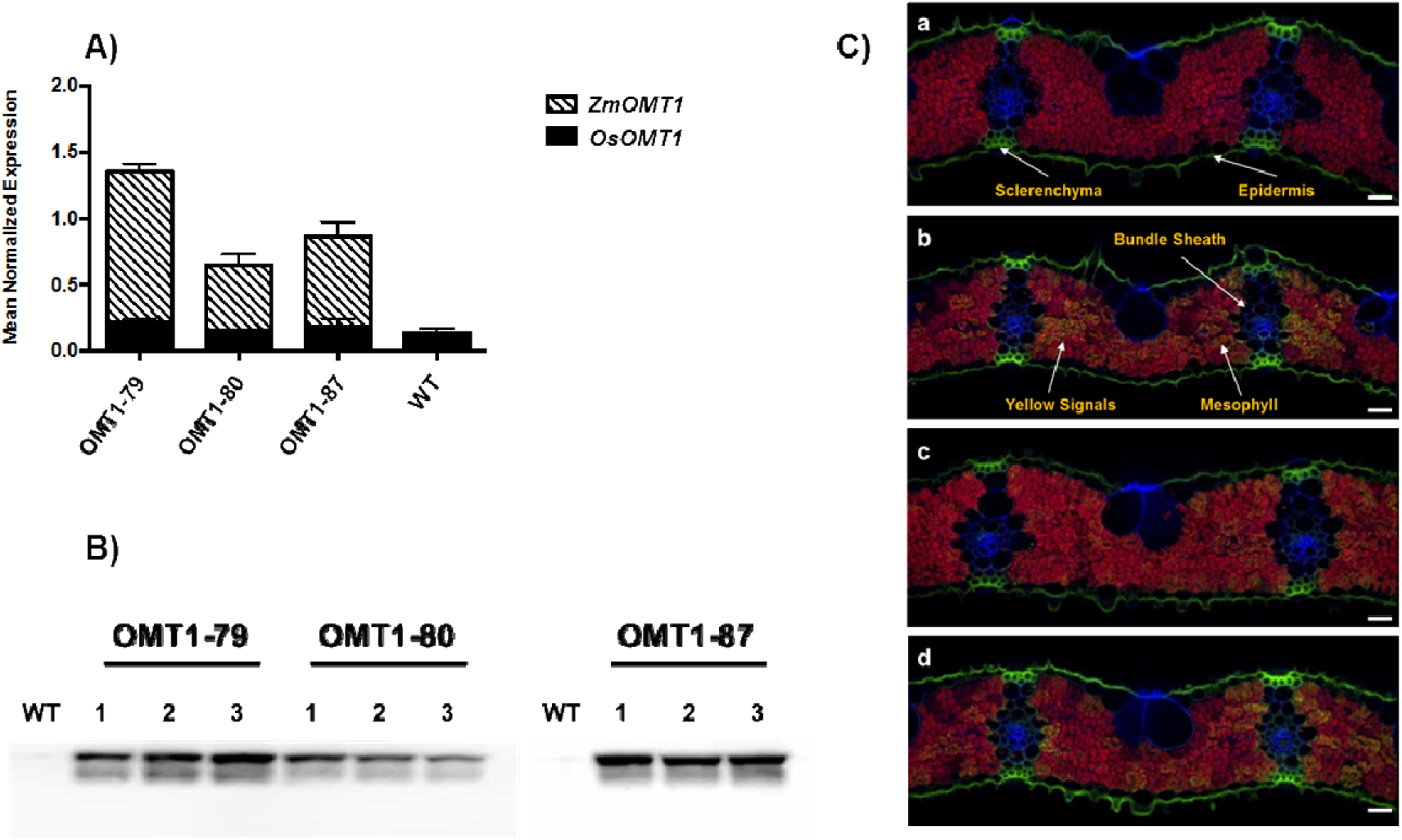
Transcript accumulation of *OsOMT1* and *ZmOMT1* genes via quantitative real time PCR analysis in the leaf blade of transgenic rice OMT1 lines (OMT1-79, OMT1-80 and OMT1-87) together with wild-type plants (WT) as a control. Data represent the mean normalized expression ± SEM of three and two biological and technical replicates, respectively (**A)**. Assessment of expressed *Zm*OMT1 protein in transgenic lines (OMT1-79, OMT1-80 and OMT1-87) together with wild-type rice (WT) as a control via Western-blot using 12% SDS-PAGE and two-step antibody immunodetection with 5 seconds exposure time **(B)**. Immunolocalization of *Zm*OMT1 protein in leaves of (A) wild-type, (B) OMT1-79, (C) OMT1-80 and (D) OMT1-87 plants in which the yellow signal is easiest to see in line OMT1-79 where the levels are highest. Anti-AcV5 tag primary antibody diluted 1:25 plus Alexa Fluor 488 (fluorescent dye) goat anti-mouse IgG secondary antibody diluted 1:200 was used to probe for AcV5 tag (shown in green color). Chlorophylls are shown as a red autofluorescence. The cell wall was co-stained with calcofluor white and is shown in blue. Magnification: 200x. Scale bar: 20 µm **(C)**.

### PCR screening

Transgenic plants were screened by genomic PCR to obtain homozygous transgenic lines. Leaf samples were harvested 7 days after transplanting into soil. PCR amplification was performed using the KAPA 3G plant PCR kit (Kapa Biosystem, USA). The scraped leaf tissue was directly used as the template for PCR amplification in a total volume of 10 µl employing the gene-specific primers; Fwd_primer 5’-CGTGGGATACCCTTACATGG and Rev_primer 5’-CCCGATTATCTTCCACCAGA. PCR conditions were: 95°C for 5 min, 32 cycles of 95°C for 20 sec, 60°C for 15 sec and 72°C for 30 sec; and 72°C for 1 min. The plasmid DNA was used as a positive control and non-transgenic rice leaf tissue or water were used as negative controls.

### Quantitative real time PCR (qRT-PCR)

qRT-PCR was performed to quantify the level of transcript expression for both *OsOMT1* and *ZmOMT1* genes in 8-week-old plants. RNA was extracted from leaf materials comprising 3 biological replicates using QIAGEN RNeasy Mini Kit. The one-step cDNA synthesis was carried out using LunaScript^™^ RT SuperMix Kit (NEB Biolabs, USA) followed by qRT-PCR master mix preparation using Luna® Universal qPCR Master Mix (NEB Biolabs, USA) which was finally performed in 7500 Fast Real-Time PCR System (Invitrogen, USA) in a total volume of 20 µl. PCR was guided using primer pairs as follows: OsOMT1_Fwd 5’-ATGGAATTGGGTCTGCTCCTG, OsOMT1_Rev 5’-AATCCATACCCCCACCACTG, *ZmOMT1*_Fwd 5’-GTGGGGCTATGGGTTTGTCA and *ZmOMT1*_Rev 5’-TATCTTCCACCAGAAGCCGC. The PCR conditions were: 95°C for 60 sec, 40 cycles of 95°C for 15 sec and 60°C for 30 sec followed by the measurement of the melting curve after 40 cycles for primer specificity. The primer efficiency was calculated as described by Udvardi *et al*., (2008) using different dilutions of cDNA together with the a highly stable housekeeping gene from rice, *OseEF-1a* (Os03g0177500) that was identified in a previous experiment by Jain *et al*., (2006). The mean normalized expression (MNE) for calculation of average CT was used as described by Simon (2003).

### Real-time PCR (RT-PCR)

To detect the mRNA expression of *ZmOMT1* and *ZmDiT2* genes in OMT1/DiT2 double cross lines, RT-PCR analysis was performed in 8 week-old plants. RNA was extracted from leaf materials using TRIzol reagent (Invitrogen, USA) and treated with DNase (Promega, USA). A 1 µg RNA was used to synthesize cDNA using a first-stand cDNA synthesis kit (Roche Diagnostics, Switzerland). The cDNA was normalized to 100 ng µl^−1^ and used for PCR analysis in a 10 µl reaction with gene-specific primers (5‘-CGTGGGATACCCTTACATGG and 5‘-CCCGATTATCTTCCACCAGA for *ZmOMT1*, 5‘-GTTGGAATGGCAGGACAACT and 5‘-ACCCAGCCTGAAAACATCTG for *ZmDiT2*, 5‘-CAACATTGTGGTCATTGGCC and 5‘-GCAGTAGTACTTGGTGGTCT for *OseEF-1a*). OseEF-1a was used as a positive and quality control. The PCR condition were as follows: pre-denaturation for 3 min at 95°C; 40 cycles of the polymerization reaction consisting of a denaturation step for 20 sec at 95°C, for 30 sec at 55°C and an extension step for 45 sec 72°C; and a final extension step for 3 min at 72°C.

### Leaf chlorophyll content and plant growth analysis

The upper fully expanded leaves of rice plants at mid-tillering stage (50-60 days old) were used to determine leaf chlorophyll content using the SPAD chlorophyll meter (Konica Minolta, Japan). The plant height and tiller number were measured at booting stage. The plant height was measured from soil level to the base of the flag leaf on the main tiller.

### Western blot and immunodetection of recombinant protein (ZmOMT1)

The presence of the AcV5-tagged *Zm*OMT1 protein in leaf membrane extracts of rice lines overexpressing *ZmOMT1* was checked by fractionating the isolated protein on 12% SDS-PAGE gel, followed by Western-blot analysis. Primary mouse anti-AcV5 tag 1:2,000 (Abcam plc, UK) and peroxidase-conjugated secondary (Goat anti-Mouse IgG (H+L) HRP, 1:2,500, ThermoFisher Scientific, Germany) antibodies were used for the detection of the AcV5 tag. Visualization of the stained protein on nitrocellulose membranes was carried out by a LAS-4000 Mini luminescence image analyzer (GE Healthcare, Germany) using the ECL Western Blotting Detection Reagents (GE Healthcare, Germany).

### DNA blot analysis

Genomic DNA was extracted from the leaves of mature rice plants using the potassium acetate method as described by Guillemaut and Maréchal-Drouard, (1992). A total of 16 µg genomic DNA was digested with HindIII restriction endonuclease (NEB Biolabs, UAS). Digested DNA was separated by electrophoresis on a 0.8 % agarose gel and then transferred onto Hybond N+ membrane (GE Healthcare, UK). Blots were hybridized with a digoxygenin (DIG)-labeled *ZmPEPC* promoter-specific probe synthesized using primers (Fwd 5’-TCCCGAGTTCCTAACCACAG; and Rev 5’-GTGGCTGAGGCTTCTTTTTG) and the PCR DIG Probe Synthesis Kit (Roche Diagnostics, Switzerland). The signals were detected by CDP-Star (Roche Diagnostics, Switzerland) following the manufacturer’s instructions.

### Immunolocalization

The seventh leaf at the mid-tillering stage was fixed and prepared for immunolocalization analysis as described in Lin *et al.*, (2016). The fixed leaf sections were probed with the anti-AcV5 mouse monoclonal antibody (Abcam plc, UK) diluted 1:25 in blocking solution. Alexa Fluor 488 (fluorescent dye) goat anti-mouse IgG (Invitrogen, USA) secondary antibody was used for detection and sections were examined on a BX61 using the Disk Scanning Unit attachment microscope (Olympus, Japan) with fluorescence functions.

### Total leaf membrane protein isolation

The fully expanded 3^rd^ leaf of rice at the mid-tillering stage was used for protein extraction. Leaves were homogenized to a fine powder using a nitrogen-cooled mortar and pestle. The powder was used as starting materials and total leaf membrane protein was isolated using an extraction buffer consisting of 250 mM Tris (HCl, pH=8.5), 25 mM EDTA, 30 % (w/v) sucrose, 5 mM DTT, and appropriate protease inhibitors. Two subsequent centrifugation steps at 10,000 g and 100,000 g were then performed, using a bench top centrifuge and ultra-centrifuge, respectively. Ultimately, the isolated membrane was re-suspended in 50 mM HEPES (KOH, pH 7.5), 5 mM EDTA, 2 mM DTT together with protease inhibitors (*for detailed procedure see* Furbank *et al*., 2001 and Roell *et al*., 2017). Finally, the protein concentration was measured utilizing the Pierce BCA Protein Assay Kit (ThermoFisher Scientific, Germany) following the manufacturer’s instructions.

### Reconstitution of total leaf membrane into liposomes

*In-vitro* analysis of transporter activity was carried out using a freeze-thaw-sonication reconstitution procedure in concert with forward exchange of the substrate (Palmieri *et al*., 1995). Following reconstitution, the proteoliposomes were preloaded with unlabeled malic acid to a final concentration of 30 mM (pH=7.5). Reconstituted proteins were separated from the non-reconstituted ones utilizing the size-based column chromatography technique (Sephadex G-25M columns (PD-10 column, GE Healthcare, USA) (*for detailed procedure see* Roell *et al*., 2017).

### Radioactive labeled [^14^C]-malate uptake measurement

Uptake of radiolabeled substrates in counter-exchange with non-labeled substrates was carried out during the course of one hour at six different time points (2, 4, 8, 16, 32 and 64 minutes). The reaction was started by adding 950 µl of proteoliposomes into 50 µl of [^14^C]-malate diluted in transport medium (7 mM malic acid, pH=7.5), and stopped at each of the above-mentioned time points by loading an 150 µl aliquot of the reaction mixture to an anion exchange resin column (acetate form, 100-200 mesh, Dowex AG1-X8 Resin, Bio-Rad, UAS). The resin column was previously equilibrated five times using 150 mM sodium acetate (pH=7.5). Unincorporated [^14^C]-malate was replaced by acetate in the resin column and the incorporated label was washed through a scintillation vial containing 10 ml Rotiszint® eco plus scintillation cocktail (Carl Roth, Germany). Finally, the uptake of radio-labeled substrate was measured as counts per minute (CPM) by scintillation counting. To correct for background and false positives, the entire experiment was repeated using proteoliposomes without pre-loading of the substrate of interest (*for detailed procedure see* Roell *et al*., 2017). The uptake data were further assessed relative to both internal standards and total protein content (mg) in each sample. Related graphs were made using the one-phase association equation in GraphPad Prism 6 (http://www.graphpad.com/prism/prism.htm).

### Photosynthetic CO_2_ assimilation, light response and dark respiration rates

Two individual fully expanded leaves per plant and three plants per line were measured for leaf photosynthetic CO_2_ assimilation and dark respiration rates during the tillering stage using a LI-6400XT portable photosynthesis system (LI-COR Biosciences, USA) in which a single leaf was clamped in the standard LI-COR leaf chamber. Measurements were performed on the mid-portion of the leaf blade between 08:00 h and 13:00 h at a constant airflow rate of 400 µmol s^−1^, leaf temperature of 30°C, and a leaf-to-air vapor pressure deficit of between 1.0 and 1.5 kPa. Leaves were acclimated in the cuvette for 30 min before measurements were started. The response curves of the net rate of CO_2_ assimilation (*A*, µmol CO_2_ m^−2^ s^−1^) to changing intercellular CO_2_ concentration (*C*_*i*_, µmol CO_2_ mol^−1^) were acquired by decreasing *C*_*a*_ (CO_2_ concentration in the cuvette) from 2,000 to 20 µmol CO_2_ mol^−1^ at a photosynthetic photon flux density (PPFD) of 2,000 µmol photons m^−2^ s^−1^. The 2 % oxygen entering the cuvette was set by mixing different concentration of nitrogen and oxygen in the CO_2_ free airstream through two mass flow controllers (model GFC17, Aalborg Mass Flow Systems, USA) at a flow rate of 1.5 ml m^−1^. Maximum Rubisco activity (V*cmax*) and maximum electron transport activity (J*max*) were determined using the PsFit Model (Bernacchi *et al*., 2001, 2003; Farquhar *et al*., 1980). The light-response curves were measured by increasing the PPFD from 20 to 2,000 µmol photons m^−2^ s^−1^ at a C_a_ of 400 µmol CO_2_ mol^−1^. The carboxylation efficiency (CE, µmol CO_2_ m^−2^ s^−1^ µmol CO_2_ mol^−1^), CO_2_ compensation point (Γ, µmol CO_2_ m^−2^ s^−1^) and quantum yield (□, mol CO_2_ mol^−1^ photons) were calculated as described by Lin *et al*., (2016). The dark respiration rate (*R*_*d*_) measurements were made on leaves in darkness following an acclimation at a photosynthetic photon flux density (PPFD) of 1,000 µmol photons m^−2^ s^−1^ for 10 min at a *C*_*a*_ of 400 µmol CO_2_ mol^−1^. The dark respiration rate (*R*_*d*_, µmole CO_2_ m^−2^ s^−1^) was calculated over a period of 1100-1200 s in the dark.

### Leaf gas exchange and photosynthetic measurement in tandem with the metabolite analysis

To normalize metabolite pool sizes by photosynthetic flux, two sets of rice plants of different ages (Set one: 30-35 days old and Set two: 50-55 days old) were analyzed. In order to measure photosynthetic CO_2_ assimilation and collect the samples for metabolite analysis under steady-state conditions, a custom gas exchange chamber was interfaced with a LI-COR 6400XT portable photosynthesis system (LI-COR Biosciences, USA) (Fig. S1B). The custom gas exchange chamber encased the leaf to be measured within a low-gas permeable sausage casing (5 cm diameter Nalophan, Kalle GmbH, Germany) to allow for rapid freeze-quenching of the sample. The chamber was constructed using two stainless-steel pipe sections fitted with Swagelok connections to the LI-COR sample line, one of which was capped on the end with a welded end cap. Prior to each measurement, a ~20 cm section of sausage casing was positioned between the pipe sections and sealed to the outside of the pipe sections using a small amount of silicone vacuum grease. The proximal end of a leaf blade was then sandwiched between two halves of a silicone stopper and inserted into the open pipe section with the adaxial side up. Actinic light was delivered via an LED ring light (Model R300, F&N Lighting, USA) which allowed constant, homogenous illumination of the leaf surface. Metabolic activity was rapidly quenched by freeze clamping the leaves with a liquid nitrogen-cooled copper disk attached to an aluminum handle. Fully expanded leaves of different tillers from five biological replicates were measured in the LICOR 6400XT that was attached to the sausage chamber. Flow through in the custom chamber was maintained at 700 μmol s^−1^, light intensity at 500 μmol photons m^−2^ s^−1^ and CO_2_ concentration was set to 200, 400 or 1,000 µmol CO_2_ mol^−1^. Leaf surface area was determined by taking a photograph and analyzing in ImageJ v1.51m9 (Schneider *et al*., 2012). Leaf temperature was not controlled but ranged between 25-27°C as determined from energy balance calculations. Leaves were sealed within the chamber until steady-state conditions were reached (as determined from a constant net CO_2_ fixation rate) and gas exchange measurements logged. After logging gas exchange data, the liquid nitrogen-cooled piston was inserted rapidly through the ring light onto the leaf and onto a plastic anvil, and then transferred rapidly to an aluminum-foil pouch and into liquid nitrogen. To avoid potential diurnal artifacts, all measurements (genotypes and CO_2_ treatments) were randomized and performed only during the peak photosynthetic activity of the rice plants between 9:00 am to 3:00 pm.

### Metabolite analysis (GC/MS)

The GC/MS-based metabolite measurements were performed as described by Fiehn, (2007), using ribitol as an internal standard. Leaf samples were collected by rapid freeze-quenching from the custom gas exchange chamber describe above. Freeze-quenched tissue was ground into a fine powder in liquid nitrogen using a mortar and pestle. Extracted metabolites were injected into a gas chromatograph (Agilent 7890B GC System, Agilent Technologies, USA) that was in line with a mass spectrometer (Agilent 7200 Accurate-Mass Q-TOF GC/MS, Agilent Technologies, USA). Metabolite peaks were evaluated using Mass Hunter Software (Agilent Technologies, UAS). The relative amount of each metabolite was calculated from the peak area, taking into account both the initial fresh weight used for extraction and the internal standard.

### Total free amino acid (FAA) content

FAA contents were measured using the Ninhydrin colorimetric method as described by Smith and Agiza, (1951), with minor changes. Briefly, 10 µl of metabolite extract together with 40 µl of methanol: water mixture (2.5:1 ratio) was added to 50 µl of 1 M citrate (NaOH, pH=5.2) and 100 µl of 1% (w/v) Ninhydrin (prepared in methanol: H_2_O, 2.5:1 ratio), and then heated to 95°C for 20 min. The solution was then transferred to a micro-well plate after a short centrifugation of 10 sec at 10,000 rpm. The total amino acid content was then measured in a Synergy HT plate reader (BioTek, Germany) at a wavelength of 550 nm. Data were adjusted based on the L-leucine standard curve and related dilution factor.

### Starch and sucrose contents

>The youngest fully expanded leaf during the tillering stage was harvested at 10:00 am and frozen immediately. Frozen leaf samples were ground in liquid nitrogen using a mortar and pestle. 50 mg of homogenized leaf powder was then extracted in 500 µl of ice-cold 0.7 M perchloric acid. For separating the soluble and insoluble fractions, the sample was centrifuged at 21,100 g for 10 min at 4°C. The insoluble fraction containing the starch was further washed five times with 1 ml of 80% (v/v) ethanol. After centrifugation, the supernatant was discarded, and the pellet was air dried and resuspended in 500 µl of water. The starch sample was gelatinized by boiling for 4 hours and hydrolyzed overnight at 37°C with 0.5 U of amyloglucosidase and 5 U of α-amylase. The starch content was measured as described in Smith and Zeeman, (2006). The soluble fraction containing sucrose was neutralized to pH=6 with neutralization buffer (2 M KOH, 0.4 M MES, 0.4 M KCl). After centrifugation at 21,100 g for 10 min at 4°C, the supernatant was transferred into a new tube and the remaining insoluble potassium perchlorate was discarded. The supernatant was assayed for sucrose content by enzymatic determination as described by Smith and Zeeman, (2006).

### Carbon: Nitrogen (C/N) ratio measurement

>The ratio of carbon to nitrogen as well as δ^*13*^C were analyzed based on leaf dry-weight (mg) of 30-day-old and 50-day-old transgenic using the ISOTOPE cube elemental analyzer connected to an Isoprime 100 isotope ratio mass spectrometer, EA-IRMS; Elemental Analyzer-Isotope ratio mass spectrometry (Elementar, Germany). The δ^*13*^C ratio is expressed as parts per thousand (‰) using the international standard of the Vienna Pee Dee Belemnite (VPDB).

### Transmission electron microscopy

>Rice seeds were germinated in petri dishes in distilled water for 4 days and then placed on a floating net in distilled water in a 19 L bin in greenhouses at the University of Toronto. Seedlings were fertilized with 1/3 strength hydroponic media at day three after transfer and then with full strength media every 4 days (Makino and Osmond, 1991). Plants were sampled from 09:30 am to 11:00 am when day length was over 11.5 h and light intensity in the unshaded greenhouse regularly exceeded 1,400 μmol photons m^−2^s^−1^. The middle section of the most recently fully expanded leaf was dissected into 2 mm pieces and prepared for transmission electron microscopy as previously described Khoshravesh *et al*., (2017). Leaf sections were fixed in 1 % glutaraldehyde, 1 % paraformaldehyde in cacodylate buffer (pH=6.9) and post-fixed in 2 % osmium tetroxide in cacodylate buffer (pH=6.9). Tissue samples were dehydrated in an ethanol series, embedded in Araldite 502 epoxy resin and sectioned at 60 nm for imaging with a Phillips 201 transmission electron microscope equipped with an Advantage HR camera system (Advanced Microscopy Techniques, USA).

### Generation of rice lines overexpressing both ZmOMT1 and ZmDiT2

To generate transgenic rice plants co-expressing *ZmOMT1* and *ZmDiT2*, homozygous *ZmOMT1* single transgenic T_2_ lines (OMT1-79, OMT1-80, and OMT1-45) were crossed with homozygous *ZmDiT2* single transgenic T_2_ lines (DiT2-27, DiT2-39 and DiT2-44) (Fig. S2A). The F1 progeny were selfed to produce segregating F2 populations that were used for all experiments reported here. The pSC110:*ZmDiT2*:AcV5 construct used for generating DiT2 lines contained the coding sequence of *ZmDiT2* (GRMZM2G40933) from *Zea mays* of the B73 variety and included an AcV5 epitope tag at the C-terminal end of the coding sequence. *ZmDiT2* was cloned using the primers: Fwd 5’-CACCATGGAGCTCCACCTCGCCAC and Rev 5’-TCAAGACCAGCCGCTCGCATCTTTCCAAGAGTACAGACCCAAAAATTTCCACCA GATG. Homozygous *ZmDiT2* lines were selected by PCR analysis and protein accumulation was determined on western blots (Fig. S2B).

## Results

### Three independent single transgene insertion lines accumulate ZmOMT1 protein in mesophyll cells

A total of 198 T_0_ plants were generated, of which 87 were positive for *ZmOMT1* as determined by PCR analysis of genomic DNA, 40 of which carried a single copy of the *ZmOMT1* transgene as determined by DNA gel blot analysis. Three single-insertion lines (OMT1-79, OMT1-80 and OMT1-87; Fig. S3) were advanced to succeeding generations to obtain homozygous lines. Homozygous plants in either the T_3_ or T_4_ generation were used for all subsequent experiments. To compare steady-state transcript levels of native rice *OsOMT1* and the introduced *ZmOMT1*, qRT-PCR was performed. Expression of the native *OsOMT1* was not affected by expression of *ZmOMT1* in any of the three over-expressing lines, with similar transcript levels observed as in wild-type rice (Fig. 1A). *ZmOMT1* transcripts accumulated in all three lines with the highest levels in OMT1-79 and the lowest in OMT1-80 (Fig. 1A). To test whether the high amounts of *ZmOMT1* mRNA in the transgenic lines was accompanied by increased transporter protein abundance, the amounts of *Zm*OMT1 protein in extracted total membrane leaf protein were examined via Western-blot, taking advantage of the C-terminal AcV5-tag. The *Zm*OMT1 protein was clearly detectable in all three lines (OMT1-79, OMT1-80 and OMT1-87) by immunoblotting (Fig. 1B). As with the transcript levels, OMT1-79 and OMT1-87 lines accumulated more *Zm*OMT1 protein than the OMT1-80 line. We further examined the spatial localization of *Zm*OMT1 in the transgenic lines by immunolocalization. Fig. 1C shows that the *Zm*OMT1 protein accumulated primarily in chloroplasts of M cells. Collectively, these data show that the *ZmPEPC* promoter drives expression of *ZmOMT1* predominantly in M cells of rice leaf tissues and that the protein can be detected in the chloroplasts of those cells.

### OMT1 membrane transporter activity is significantly increased in transgenic rice lines

To test whether expression of the *ZmOMT1* transgene led to increased OMT1 transporter activity in transgenic lines, we measured malate counter-exchange activity in liposomes reconstituted with membrane proteins isolated from wild-type and overexpressing lines (Fig. 2A). We detected significantly higher malate-malate counter-exchange activity in liposomes reconstituted with membrane proteins from overexpression lines as compared to liposomes reconstituted with membrane proteins isolated from the wild types. These data clearly indicate that the recombinantly introduced *Zm*OMT1 transporter protein is active in rice (Fig. 2B).

**Fig. 2:**
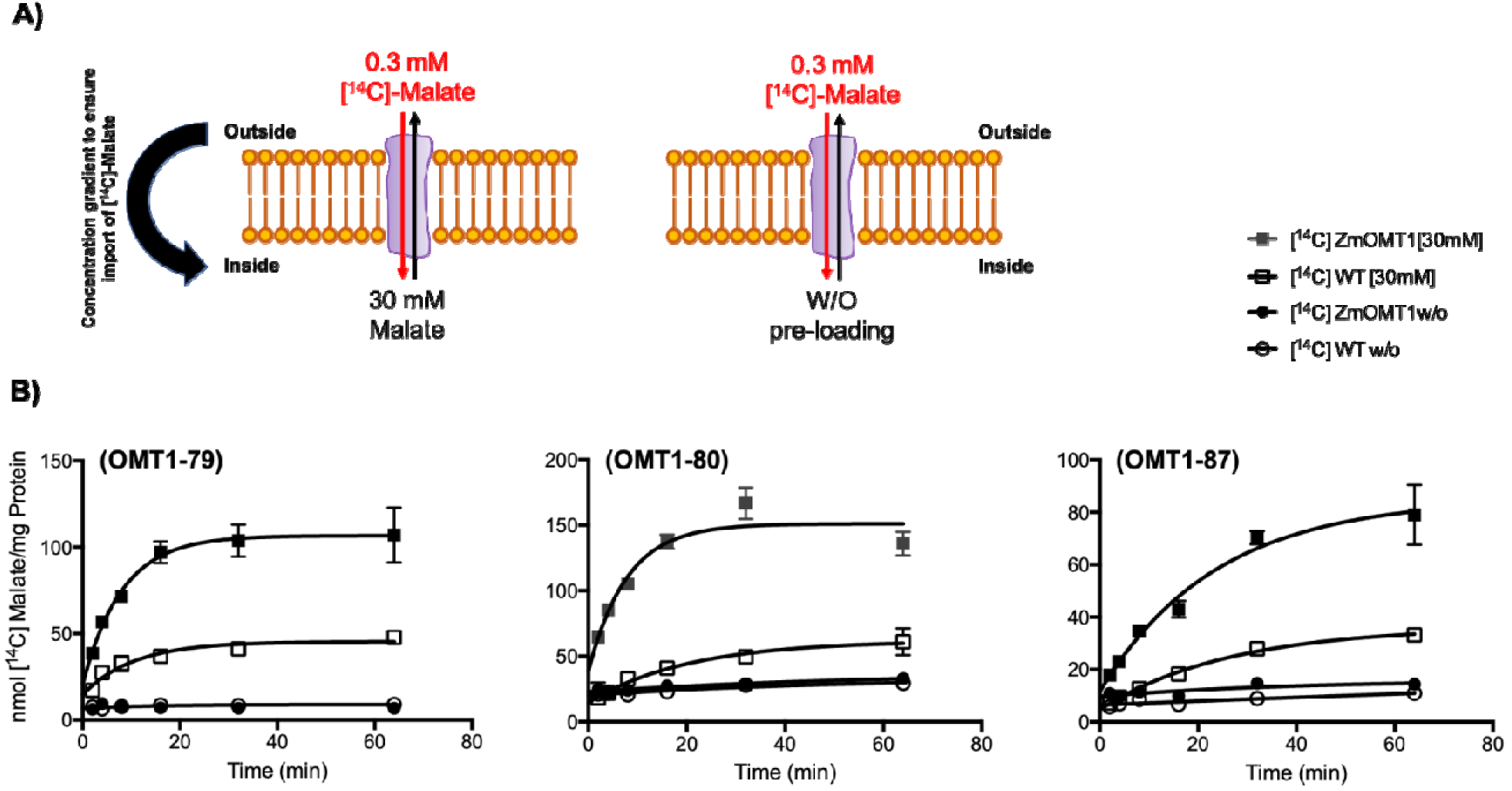
Illustration of proteoliposome after reconstitution of the total leaf membrane protein. Uptake of [^14^C]-malate was measured with 30 mM or without (w/o) pre-loading of unlabeled malate inside the proteoliposome. The activity was initiated with a final concentration of 0.3 mM [^14^C]-malate **(A)**. Uptake of malate by total crude membrane protein of wild type rice (WT) as a control together with three different transgenic OMT1 lines (OMT1-79, OMT1-80 and OMT1-87). Values represent the mean ± SEM, n=3 **(B)**.

### Slower growth and leaf lesion phenotypes of OMT1 lines

>The transgenic plants with the highest *Zm*OMT1 protein levels (OMT1-79 and OMT1-87) displayed perturbed phenotypes at the whole plant level. The OMT1-79 and OMT-87 lines were shorter (Fig. 3A and Table 1) than wild-type and displayed lesions in mature leaves in IRRI (Fig. 3B). An ELISA test for detection of infection caused by tungro virus was negative (data not shown), indicating that the lesions were not caused by tungro virus infection. The OMT1-80 line that accumulates lower levels of *Zm*OMT1 (Fig. 1) had more and longer tillers compared to wild-type (Fig. 3A and Table 1) and did not have a lesion mimic phenotype (Fig. 3B). Despite the different lesion mimic phenotypes, chlorophyll content was similar in the youngest fully expanded leaves of all three transgenic lines and wild-type (Table 1). These results suggest that high levels of *ZmOMT1* expression in rice inhibit plant growth and induce a lesion mimic phenotype in mature leaves, without altering chlorophyll content in young leaves.

**Table 1:** Leaf chlorophyll content, plant height and tiller number of wild-type and OMT1 lines.

|  | Chl (SPAD value) | Tiller number | Plant height (cm) |
| --- | --- | --- | --- |
| Wild-type | 42.1±0.7<sup>ns</sup> | 9±0.5<sup>ab</sup> | 56.4±1.1<sup>a</sup> |
| OMT1-79 | 42.4±1.4<sup>ns</sup> | 8±2<sup>b</sup> | 42.3±0.4<sup>c</sup> |
| OMT1-80 | 41.3±1<sup>ns</sup> | 12±1<sup>a</sup> | 60±1.5<sup>a</sup> |
| OMT1-87 | 42.2±0.8<sup>ns</sup> | 6.8±0.9<sup>b</sup> | 47.8±2.9<sup>b</sup> |

Chl SPAD values are the average ± SEM of three leaves from four plants at mid-tillering stage using the upper fully expanded leaves.
Tiller number and plant height are the average ± SEM of four individual T<sub>3</sub> plants; 70 days post-germination.
Different letters within groups indicate that values are statistically different $p \leq 0.05$ , Tukey's multiple comparison test. ns indicates non-significant, $p > 0.05$ .

**Fig. 3:**
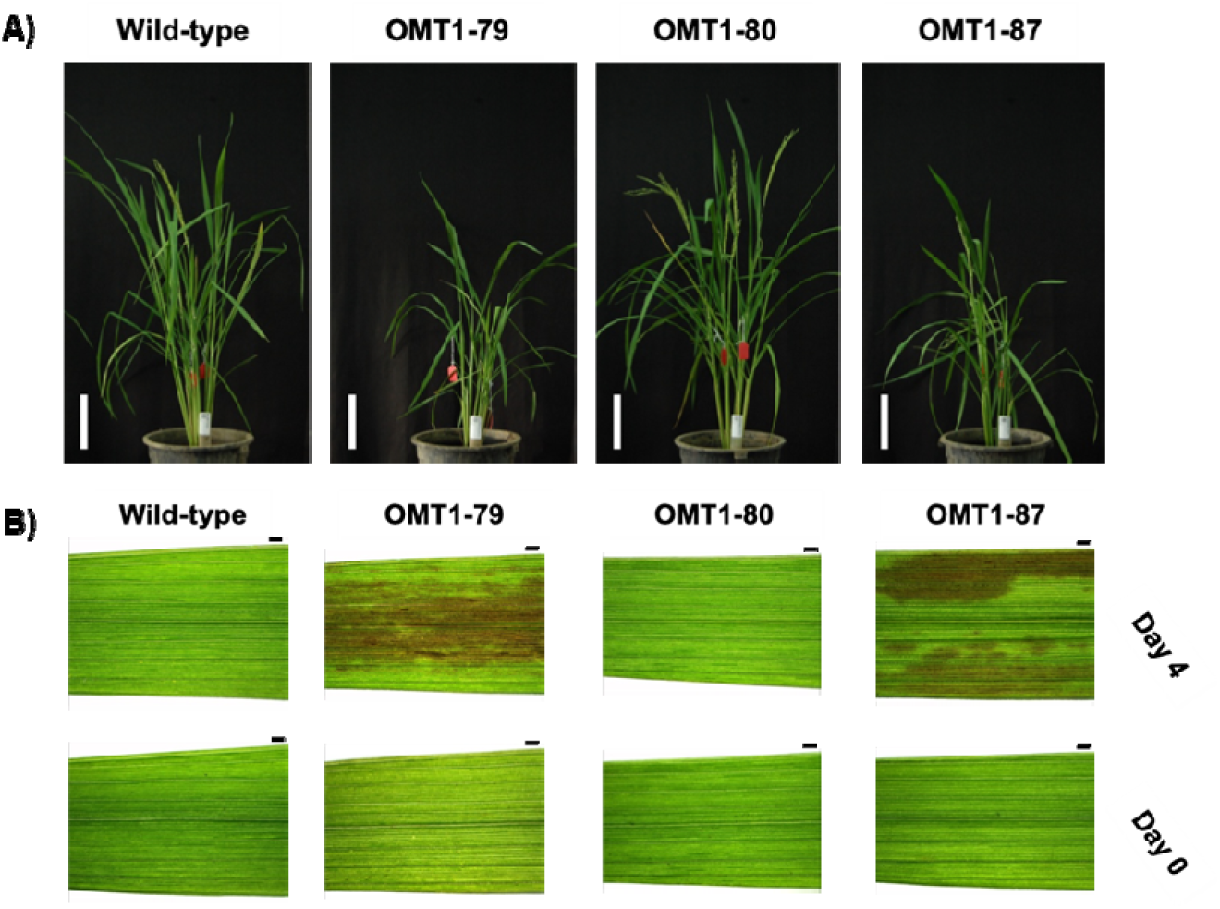
Representative pictures of wild-type, OMT1-79, OMT1-80 and OMT1-87 lines grown under ambient conditions; 70 days post germination (DPG). Scale bar: 15 cm **(A)**. Representative pictures of youngest fully expanded leaves (Day 0) and the same leaves after 4 days (Day 4) of wild-type, OMT1-79, OMT1-80 and OMT1-87 plants. The middle portions of youngest fully expanded leaves were taken when their next leaf needles started to emerge (Day 0) and the same positions from the same leaves were taken after 4 days (Day 4). Scale bar: 1 mm **(B)**.

### Photorespiratory-deficient phenotypes of ZmOMT1 transgenic lines

To examine the effect of overexpressing ZmOMT1 on photosynthesis in response to changing light conditions, the CO_2_ assimilation rate (A) in response to photosynthetic photon flux density (PPFD) was measured at ambient CO_2_ condition (400 µmol CO_2_ mol^−1^). The transgenic lines with highest *ZmOMT1* expression (OMT1-79 and OMT1-87) had slightly lower CO_2_ assimilation rates than wild-type whereas OMT1-80 had a similar rate (Fig. 4a). At 2000 µmol photon m^−2^ s^−1^, photosynthesis in the OMT-80 line and wild-type was already saturated, but this was not the case for OMT-79 and OMT-87 lines. The *ZmOMT1* over-expressing lines had similar quantum efficiency (QE) from the initial slope of light response curves (PPFD < 100 µmol photons m^−2^ s^−1^) to wild-type (Table 2) suggesting that overexpressing ZmOMT1 protein doesn’t affect the efficiency of using light energy to fix CO_2_ in rice plants. The dark respiration rates were twice as high in OMT1-79 and OMT1-87 lines compared to OMT1-80 and wild-type (Table 2) suggested that the carbon balance is possible altered in OMT1-79 and OMT1-87 compared to wild-type. Moreover, the CO_2_ assimilation rate (A) in response to intercellular CO_2_ concentration (C_i_) under non-photorespiratory (2% O_2_) versus photorespiratory (21% O_2_) conditions was measured under saturating light intensity of 2000 µmol photons m^−2^s^−1^. At 21% O_2_, lower photosynthetic rates were observed in OMT1-79 and OMT1-87 lines compared to wild-type and the OMT1-80 line (Fig. 4B). OMT1-79 and OMT1-87 lines also had higher CO_2_ compensation points (Γ) and lower carboxylation efficiencies (CE) (Table 2). Under low photorespiratory conditions (2% O_2_), wild-type, OMT1-80, and OMT1-87 had similar photosynthetic rates at ~ 40 µmol CO_2_ mol^−1^, and similar CO_2_ compensation points (Γ) (Fig. 4C). Above a C_i_ of 400 µmol CO_2_ mol^−1^, the assimilation rate was lower in OMT1-79 and OMT1-87 lines. The maximum rate of Rubisco carboxylation (Vcmax) and the maximum rate of electron transport (Jmax) were reduced in OMT1-79 and OMT1-87 lines under high photorespiratory conditions (Table S1). Together, these results indicate that the transgenic lines are Rubisco-limited under high photorespiratory conditions (21% O_2_) and that when *ZmOMT1* is expressed, ribulose 1,5-bisphosphate (RuBP) regeneration is limited at high CO_2_ concentrations. Together, these data indicate that *ZmOMT1* over-expression lines leads to higher rates of photorespiration, a suggestion supported by the observation that transgenic lines have a higher CO_2_ compensation point (Γ) than wild-type at 21% but not at 2 % O_2_.

**Table 2:**
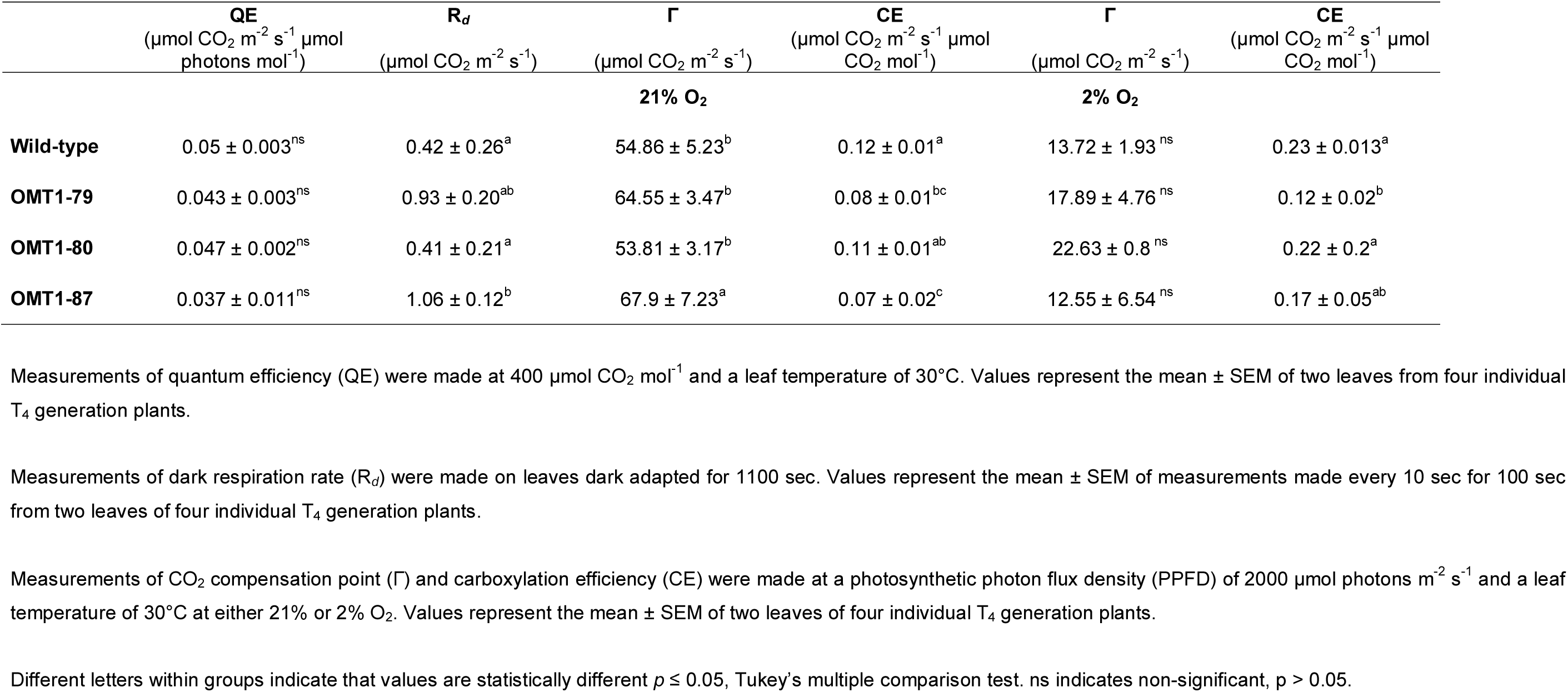
Comparison of photosynthesis parameters.

**Fig. 4:**
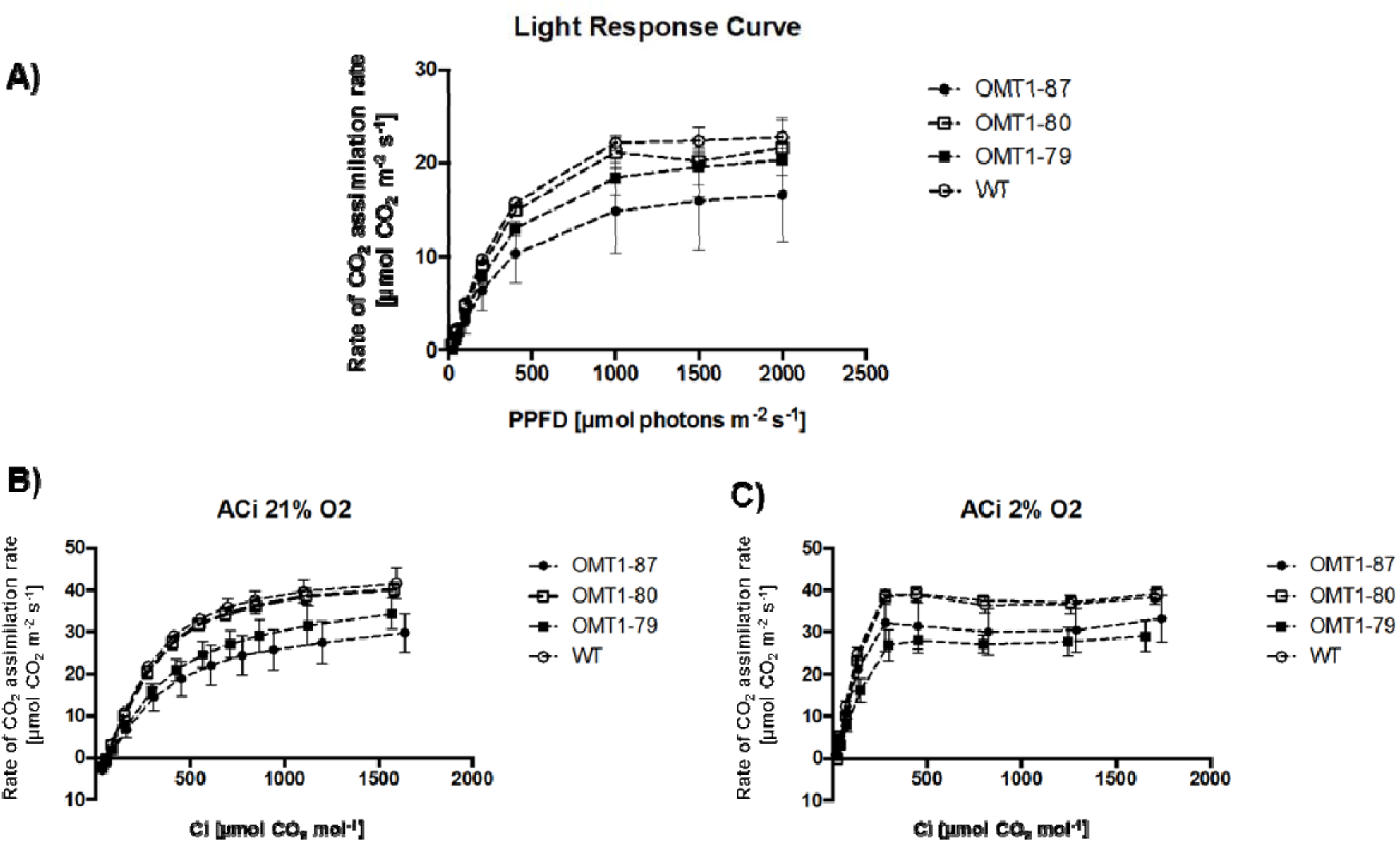
Rate of CO_2_ assimilation in response to photosynthetic photon flux density (PPFD). Light response curve measurements were carried out under 400 µmol CO_2_ mol^−1^ and leaf temperature of 30°C. Values represent the mean ± SEM of two leaves of four individual T_4_ plants of OMT1 lines (OMT1-79, OMT1-80 and OMT1-87) and wild-type rice (WT) **(A)**. Rate of CO_2_ assimilation in response to intercellular CO_2_ concentration (Ci) at 21% **(B)** and 2% O_2_ (**C**). The measurements were carried out under the light intensity of 2000 µmol photons m^−2^ s^−1^ with the leaf temperature of 30°C. Values represent the mean ± SEM of two leaves of four individual T4 plants of OMT1 lines (OMT1-79, OMT1-80 and OMT1-87) and wild-type rice (WT).

### Chloroplast ultrastructure is perturbed in OMT1 transgenic lines

The macroscopic and physiological phenotypes of OMT1 lines were accompanied by ultrastructural changes in M cell chloroplasts. In contrast to wild-type plants, M cell chloroplasts of the OMT1 lines developed a peripheral reticulum (PR; Fig. 5) which is an internal network of tubules and vesicles continuous with the chloroplast inner membrane of chloroplasts (Rosado-Alberio *et al*., 1968, Laetsch, 1974). Plastoglobules (PG), not observed in wild-type plants, were also present in chloroplasts of the over-expressing lines. PGs are lipid microcompartments posited to function in lipid metabolism, redox and photosynthetic regulation and thylakoid repair and disposal during chloroplast biogenesis and stress (Rottet *et al*., 2015; van Wijk and Kessler, 2017).

**Fig. 5:**
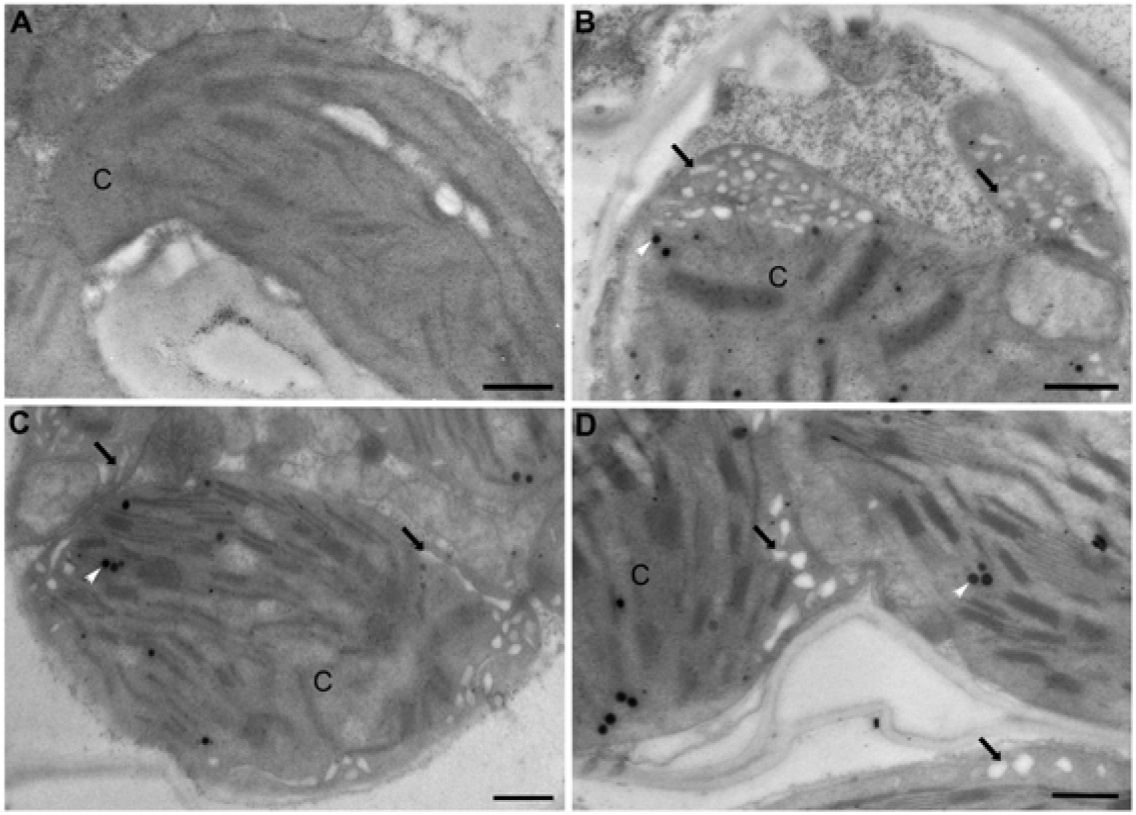
Transmission electron micrographs illustrating chloroplasts without peripheral reticulum in **(A)** wild-type and with peripheral reticulum (black arrows) in OMT1 transgenic lines **(B)** OMT1-79, **(C)** OMT1-80 and **(D)** OMT1-87. White arrows mark plastoglobules; C, chloroplast; Scale bar = 500 nm.

### CO_2_ assimilation rate and leaf metabolite profiles of transgenic lines

The photosynthetic rate of the older *ZmOMT1* transgenic plants (50-55 days old) measured in our custom-build gas exchange cuvette (Fig. S1B) was affected more than that of younger ones (30-35 days old) (Fig. 6A and B), in which the photosynthesis rate became significantly lower in *ZmOMT1* transgenic lines under ambient CO_2_ concentration (400 ppm) (Fig. 6 B). The photosynthetic rate was partially restored under high CO_2_ concentration (1000 ppm) for older plants (Fig. 6B).

**Figure 6:**
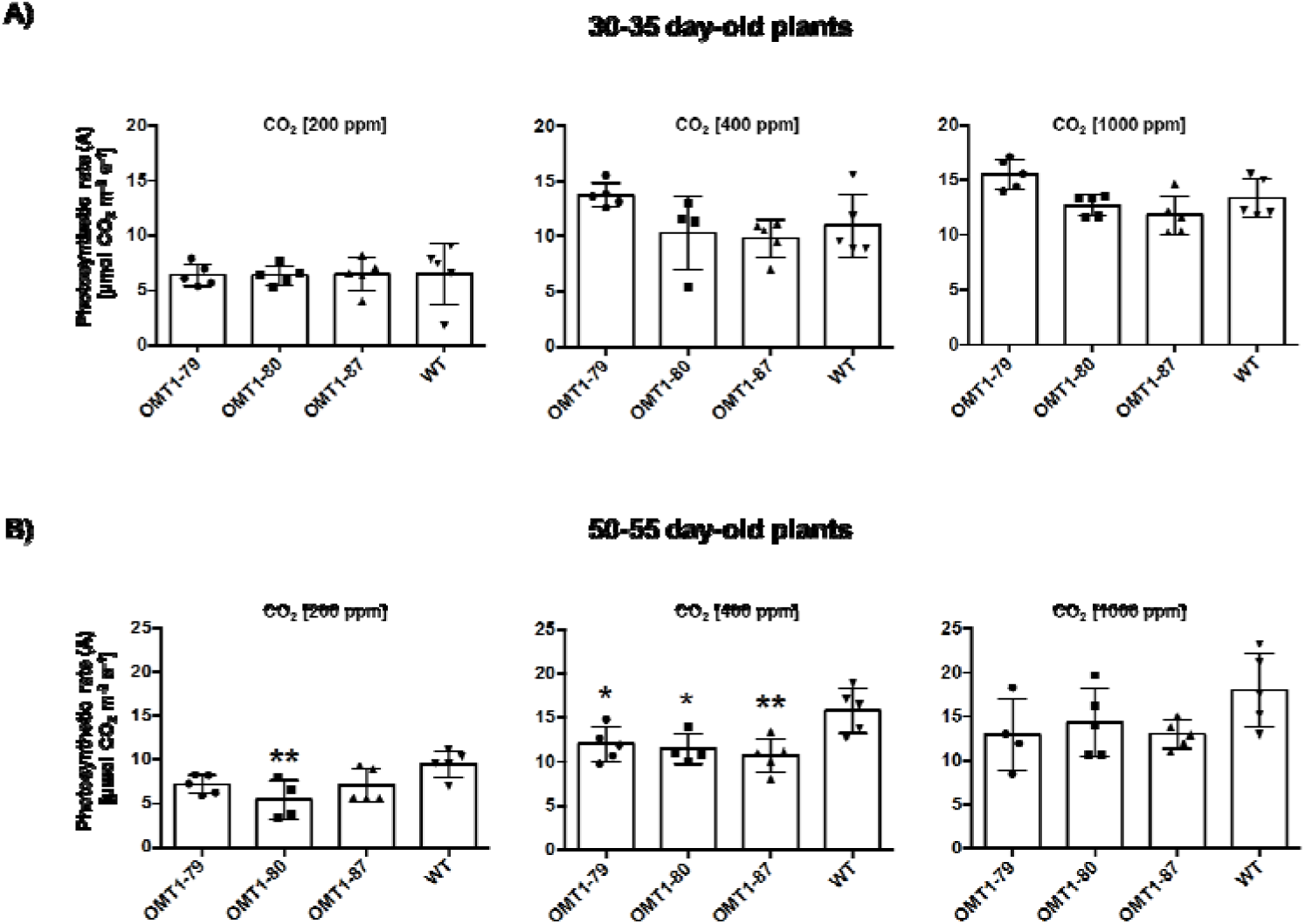
Impact of three CO_2_ concentrations (200, 400 and 1000 ppm) on photosynthetic rate measured inside a custom gas exchange cuvette of two different plants set [OMT1 lines (OMT1-79, OMT1-80 and OMT1-87) and wild-type (WT)] differed in age younger: 30-35 days old **(A)** and older: 50-55 days old **(B)**. Values represent the mean ± SEM, n=4-5. Significantly differences to WT are indicated by * *P* ≤ 0.05 and ** *P* ≤ 0.01, Tukey’s multiple comparison test.

### Metabolite profiles of ZmOMT1 lines and wild-type rice reveal altered steady state pools of TCA intermediates and aspartate

The metabolic state of 30-35-day-old *ZmOMT1* transgenic rice lines and wild-type under different CO_2_ conditions was examined using GC/MS analysis. Large differences were observed among the measured metabolites of the mitochondrial tricarboxylic acid cycle between the transgenic lines and wild-type. Malic acid, fumaric acid, iso-citric acid, succinic acid, and α-ketoglutarate were significantly lower in all *ZmOMT1* transgenic rice lines than wild-type under different CO_2_ concentrations (Fig. 7A). Among photorespiratory intermediates, only glyceric acid displayed a lower amount in OMT1 lines. Others, such as glycolic acid, glycine, and serine were similar to the wild type or tended to be higher, in some cases significantly (Fig. 7B). Of the substrates transported by OMT1 and DiT2, apparently, aspartic acid was significantly increased in the overexpression lines (Fig. 7C). Malic acid and α-ketoglutarate, as previously mentioned, were significantly lower and glutamic acid remained unchanged for all three OMT1 transgenic rice lines in comparison with wild-type under different CO_2_ concentrations (Fig. 7A and 7C). We further calculated the aspartate/malate ratio for all transgenic rice lines and compared to wild-type. As shown in Figure 7D, the aspartate to malate ratio was significantly higher in transgenic *ZmOMT1* lines relative to wild-type under different CO_2_ concentrations.

**Fig. 7:**
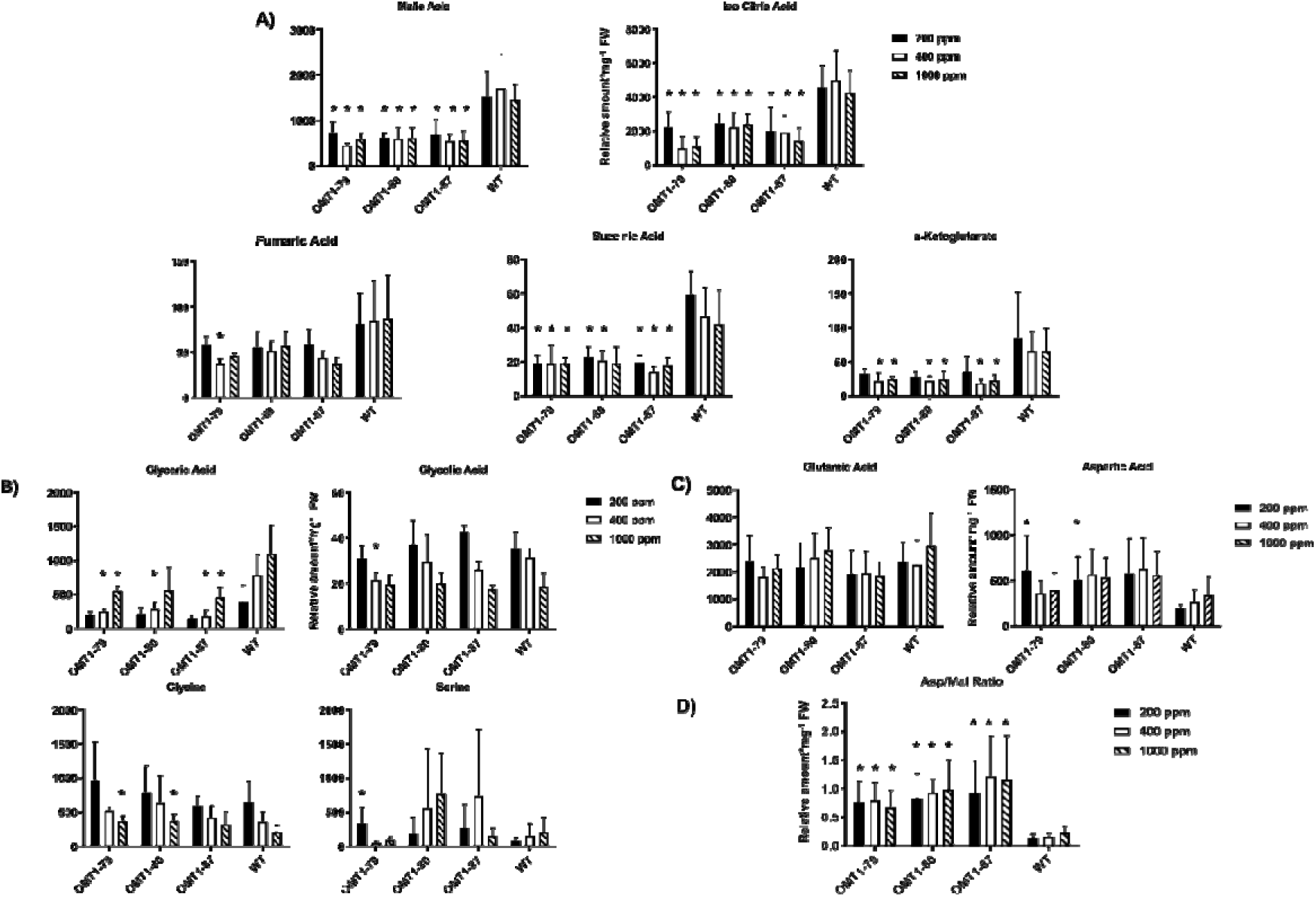
Relative amount of metabolites involved in the citric acid cycle extracted from OMT1 lines (OMT1-79, OMT1-80 and OMT1-87) and wild-type (WT) rice leaves under different CO_2_ concentrations (200, 400 and 1000 ppm). Values represent the mean ± SEM, n=5, significantly differences to WT are indicated by * *P* ≤ 0.05, Student’s t-test **(A)**. Relative amount of metabolites related to photorespiration extracted from OMT1 lines (OMT1-79, OMT1-80 and OMT1-87) and wild-type (WT) rice leaves under different CO_2_ concentrations (200, 400 and 1000 ppm). Values represent the mean ± SEM, n=5, significantly differences to WT are indicated by * *P* ≤ 0.05, Student’s t-test **(B)**.Relative amount of metabolites known as the key substrates of OMT1 and DiT2 membrane transporters extracted from OMT1 lines (OMT1-79, OMT1-80 and OMT1-87) and wild-type (WT) rice leaves under different CO_2_ concentrations (200, 400 and 1000 ppm). Values represent the mean ± SEM, n=5, significantly differences to WT are indicated by * *P* ≤ 0.05, Student’s t-test **(C)**. Aspartate/malate ratio of OMT1 lines (OMT1-79, OMT1-80 and OMT1-87) and wild-type (WT) rice under different CO_2_ concentrations (200, 400 and 1000 ppm). Values represent the mean ± SEM, n=5, significantly differences to WT are indicated by * *P* ≤ 0.05. Student’s t-test **(D)**.

### Total free amino acids, carbon:nitrogen ratios, and carbohydrate contents are decreased in leaves of ZmOMT1 lines

The absolute FAA contents of *ZmOMT1* lines and wild-type rice were determined to assess the effect of altered plastidial dicarboxylate transport capacity on amino acid metabolism. Levels were lower in older plants of *ZmOMT1* lines (50-55 days old) under all CO_2_ concentrations but were significantly decreased under ambient CO_2_ (400 ppm) compared to wild-type rice (Fig. 8A). As plants aged, the C/N ratio also decreased significantly in *ZmOMT1* transgenic lines but the δ*13*C value did not differ between wild-type and transgenic lines (Fig. 8B). Sucrose and starch amounts were significantly reduced in the OMT1 lines compared to wild-type plants (Fig. 8c).

**Figure 8:**
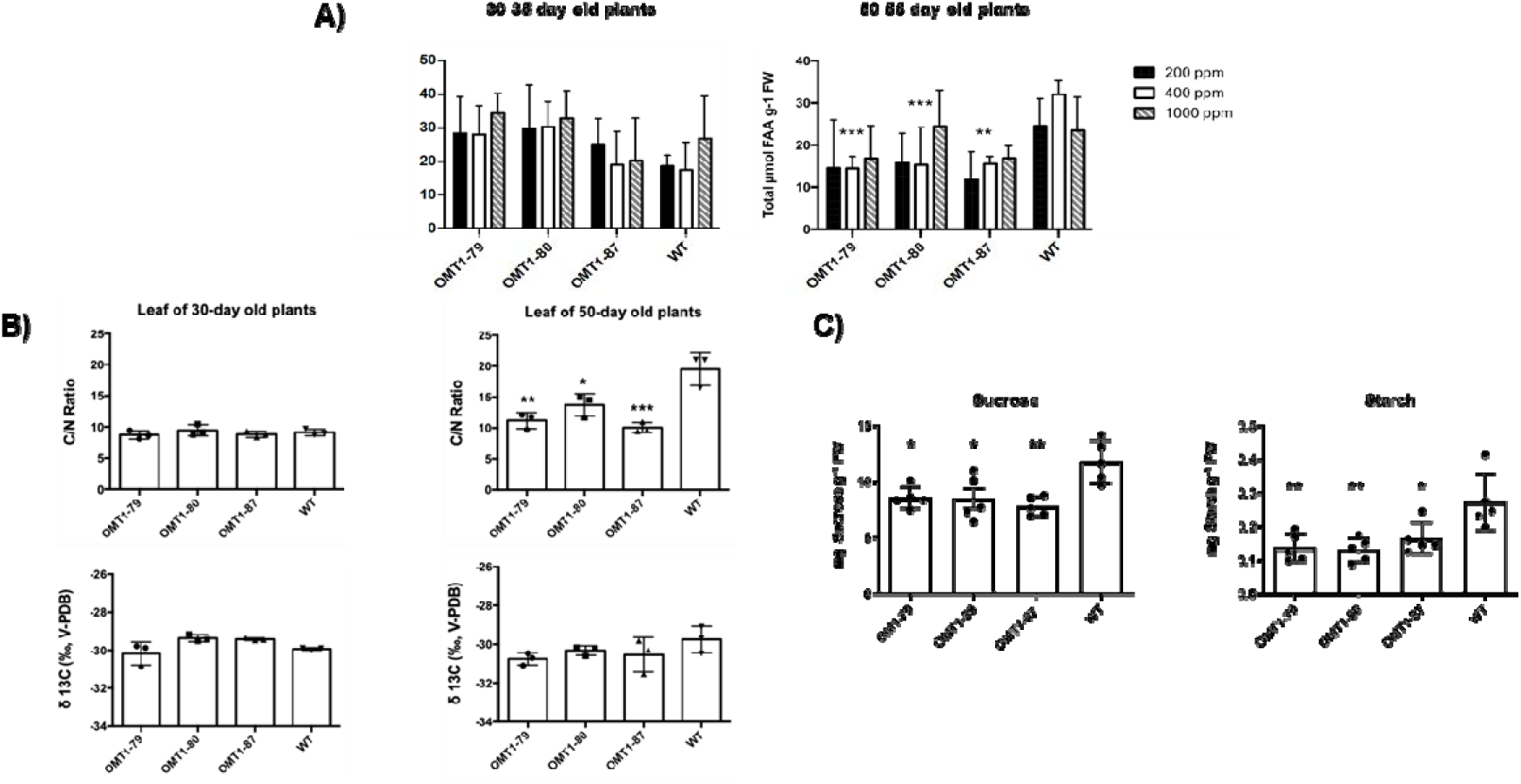
Absolute amounts of total free amino acid (FAA) content of OMT1 lines (OMT1-79, OMT1-80 and OMT1-87) and wild-type (WT) rice under different CO_2_ concentrations (200, 400 and 1000 ppm). Values represent the mean ± SEM, n=5, significantly differences to WT are indicated by ** *P* ≤ 0.01 and *** *P* ≤ 0. 001, Tukey’s multiple comparison test **(A)**. The C/N ratio and δ^*13*^C value of 30 and 50 days old OMT1 lines and wild-type rice (WT). Values represent the mean ± SEM, n=3, significantly differences to WT are indicated by * *P* ≤ 0.05, ** *P* ≤ 0.01 and *** *P* ≤ 0. 001, Tukey’s multiple comparison test **(B)**. Sucrose and starch content in OMT1 lines (79, 80 and 87) and wild-type rice (WT). Sample materials were collected at 10:00 h at the mid-tillering stage. Values represent the mean ± SEM, n=5, significantly differences to WT are indicated by * *P* ≤ 0.05, and ** *P* ≤ 0.01, Tukey’s multiple comparison test **(C)**.

### Simultaneous expression of ZmOMT1 and ZmDiT2 in transgenic rice lines restored the wild-type growth phenotype

We hypothesized that the phenotypes observed in rice lines overexpressing *Zm*OMT1 might be caused by an imbalance between the transport capacities for malate, oxaloacetate, and α-ketoglutarate (transported by OMT1), and glutamate and aspartate (transported by DiT2). If this assumption was true, then the phenotypes of *ZmOMT1* single transgenic lines should be rescued by simultaneous overexpression of *ZmDiT2*. We, hence, generated double transgenic lines in which both, *ZmOMT1* and *ZmDiT2* were expressed (Fig. S5 and S6). Notably, double transgenic lines displayed similar physiological phenotypes as wild-type plants when grown under ambient conditions (Fig. 9). Leaf chlorophyll content, number of tillers, and plant height were comparable to wild-type in two of three independent *ZmOMT1/ZmDiT2* double over-expressing plants.

**Fig. 9:**
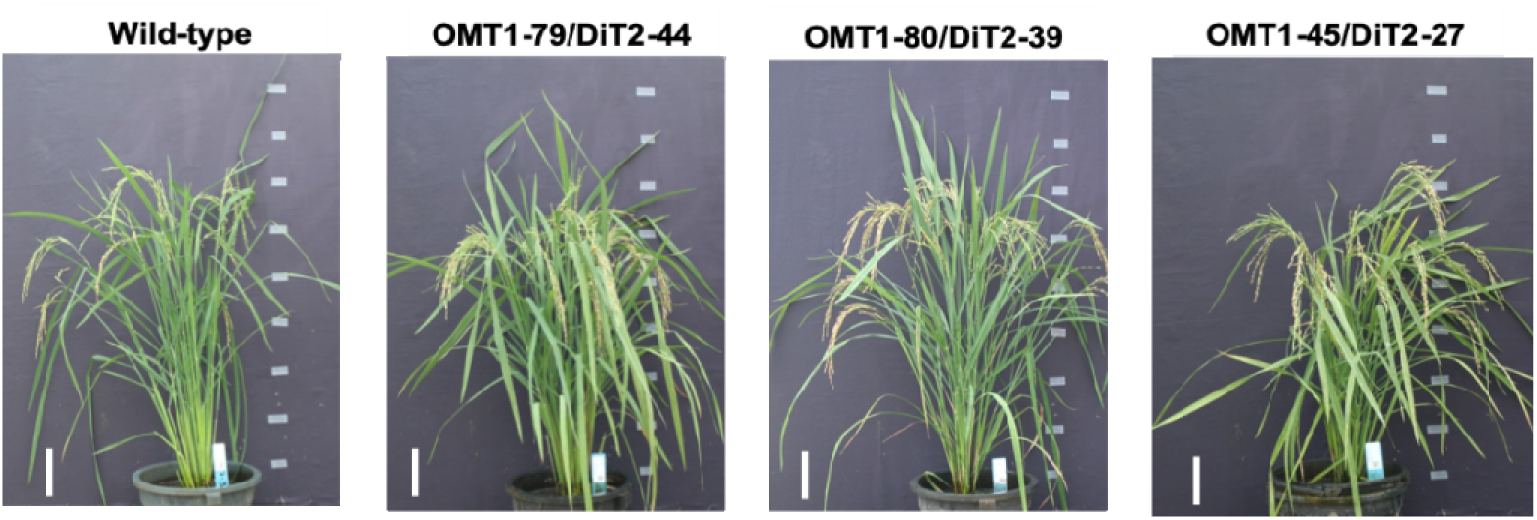
Growth phenotype of wild-type together with the double over-expressed lines of OMT1-79/DiT2-44, OMT1-80/DiT2-39, and OMT1-45/DiT2-27. All plants were grown under ambient conditions; 90 days post germination (DPG). Scale bar: 10 cm.

## Discussion

C_4_ plants require a higher transport capacity of oxaloacetate and malate across the chloroplast envelope of leaf M cells because OAA generated by the PEP carboxylase reaction in the cytoplasm is further converted to malate by plastidial NADP-malate dehydrogenase. Malate is then exported from M chloroplasts and transported to the carbon concentrating sheath cells. In this study, as part of the effort to engineer C_4_ rice, transgenic rice lines were generated that over-express the chloroplast envelope malate/oxaloacetate/α-ketoglutarate antiporter OMT1 from maize, *ZmOMT1*.

A striking feature of chloroplasts in the *ZmOMT1* overexpressing lines was the development of the peripheral reticulum (PR). This peripheral matrix of tubules and vesicles is continuous with the inner envelope which is the site where metabolite exchange occurs (Pottosin and Shabala, 2016). Although PR has been reported to be present in M and bundle sheath cells of other C_3_ grasses such as wheat (Szczepanik and Sowinski, 2014), this cellular feature has not been observed in other *Oryza* species or cultivars (Sage and Sage, 2009; Giuliani *et al*., 2013). The PR is also present in M and sheath cells of C_4_ species of grasses and eudicots, although in comparison to C_3_ grasses, the PR in C_4_ species is much more abundant (Rosado-Alberio *et al*., 1968; Laetsch, 1968; Laetsch, 1969; Szczepanik and Sowinski, 2014). Chloroplast envelope proliferation in association with over-expression of envelope proteins has been previously reported (Breuers *et al*., 2012), supporting the idea that the *Zm*OMT1 transporter is accumulating to high amounts in the inner envelope of M chloroplasts. Given that the presence of PR is posited to be correlated with high rates of metabolite exchange (Gracen *et al*., 1972a, Hilliard and West, 1971; Laetsch, 1974; Gracen *et al*., 1972b), the PR phenotype in *ZmOMT1* transgenic lines is consistent with the altered metabolic profiles observed.

In general, OAA transported by OMT1 enters the chloroplast and is subsequently converted to either malate by NADP-MDH or aspartate by plastidial aspartate aminotransferase. Whereas, malate can be transported back to the cytosol by OMT1, export of aspartate out of chloroplast requires the activity of DiT2. Enhanced accumulation of aspartate in the transgenic lines (Fig. 7D) indicates that this metabolite cannot be further metabolized in chloroplasts and thus that metabolite flux between chloroplasts and mitochondria is blocked. This outcome could explain the lower amounts of intermediate metabolites in the citric acid cycle (TCA) of mitochondria (the energy machinery) (Fig. 7A) among which a few are common substrates of the OMT1 transporter (Fig. 7A and 7C). All of these intermediates are pivotal for effective function of plant metabolic pathways. For instance, malate, a primary substrate of OMT1, participates as an intermediate in many vital mechanisms in the cytosol and vacuole (redox homeostasis, pH levels and carbon storage) (Fernie and Martinoia, 2009). Loss of function mutations in *OMT1* in the C_3_ plants Arabidopsis (Kinoshita *et al*., 2011) and tobacco (Schneidereit *et al*., 2006), caused an increase in levels of 2-OG and malate and a decrease in levels of aspartate, the opposite trend to that seen in *ZmOMT1* overexpressing rice plants. Surprisingly, any disruption to OMT1 activity (either an increase or decrease) leads to lower photosynthetic rates than wild-type, suggesting that OMT1 transporter activity must be precisely regulated to maintain optimal photosynthetic performance. The reduced photosynthetic rates in *ZmOMT1* transgenic rice plants reveal possible relationships between photosynthesis, photorespiration, and cellular redox status. Differences in photosynthesis were significant in the plants measured in the Philippines and in older plants grown in Düsseldorf, Germany (Fig. 4A and 6B). This decrease in photosynthesis is only partially explained by increases in R_*d*_ (Table 2). Interestingly, this decrease in photosynthesis could be rescued by minimizing photorespiration under some measurement and growth conditions, but not others. Specifically, the photosynthetic rates of *ZmOMT1* transgenic lines were not rescued by elevated CO_2_ or reduced O_2_ when measured under growth conditions in the Philippines (Fig. 4B and 4C), but were rescued in the plants grown in Düsseldorf, Germany when measured under elevated CO_2_ (Fig. 6). One major difference in these measurements was the light intensity used (2000 μmol m^−2^ s^−1^ for the A-C_i_ curves vs 500 μmol m^−2^ s^−1^ for the metabolite assays), meaning that phenotypic rescue may only occur under sub-saturating light intensities. As photorespiratory rates increase, the increased demand for ATP relative to NAD(P)H pushes the redox status of the NADP^+^/NADPH pools to be more reduced unless processes either decrease plastidic NADPH (malate valve) or increase ATP production (cyclic electron flux around photosystem I, CEF). The oxidation of NADPH, which could be increased with increased export of malate, must be finely balanced with metabolic demand so as not to directly compete with NADPH pools needed to supply the Calvin-Benson cycle or photorespiration. Under sub-saturating light, there are numerous lines of evidence suggesting that the malate valve regulates this balance, particularly under photorespiratory conditions (Kramer and Evans, 2011; Walker *et al*., 2014; Shameer et al., 2019). Specially, this event leads to the reduced provision of carbon skeletons for nitrogen assimilation and to a significant reduction of the leaf C/N ratio (Fig. 8B) together with the reduction of FAA in the older OMT1 transgenic lines under 400 ppm CO_2_ concentration (Fig. 8A). Principally, both carbohydrate and amino acid biosynthesis are relying on each other (Nunes-Nesi *et al*., 2010). Correspondingly, in all three OMT1 transgenic lines, both sucrose and starch contents were decreased significantly compared to wild-type rice (Fig. 8C). It is known that a part of the photo-assimilated carbon during the day will be partitioned and stored as starch to be used later during the night as a source of energy supply for sink tissues as well as fatty acid and amino acid biogenesis (Stitt and Zeeman, 2012). On the other hand, sucrose biosynthesis is occurring during the day (from the triose-phosphate pathway) and the night (from various enzymatic reactions involved in starch degradation) (Kunz et al., 2014). Therefore, starch and sucrose metabolisms tightly depend on each other and both are orchestrated by the amount of the fixed carbon during photosynthesis. Taken together, apparently too high or too low amounts of OMT1 protein affect the coordination of the C and N assimilation pathways.

### Concluding model

>Our results present evidences on the crucial roles of OMT1 transporter in rice plants. We suggest a hypothetical model (Fig. 10) in which aspartate accumulates in chloroplast of single OMT1 transgenic lines in comparison with wild-type rice (Fig. 7D). We propose that the accumulated aspartate impairs the flux between the inside and outside of the chloroplast causing the growth and photosynthetic deficiency phenotypes in single OMT1 transgenic lines. Our assumption is supported by the finding that providing an exit pathway for aspartate by introducing an additional plastidial transporter (ZmDiT2) suppresses the phenotype of OMT1 overexpression (Fig. 10). These double over-expressor OMT1/DiT2 lines grew similar as the wild-type and plant height along with numbers of tiller were recovered (Table 3). Our results indicate that coordinated expression of OMT1 and DiT2 is needed for engineering C_4_-rice plants.

**Table 3:**
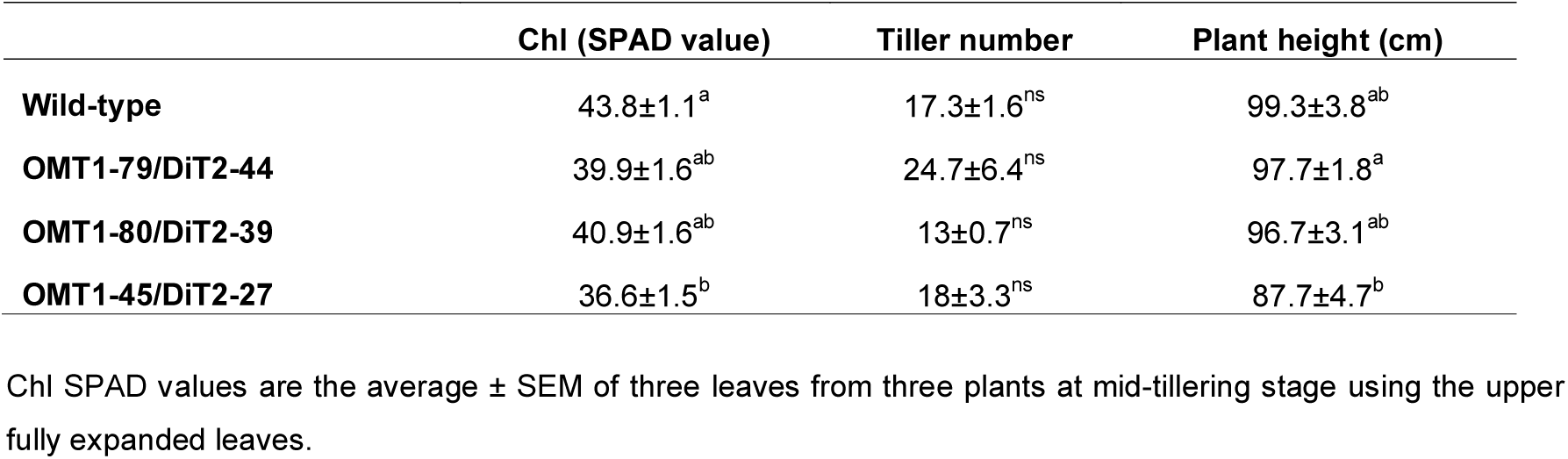
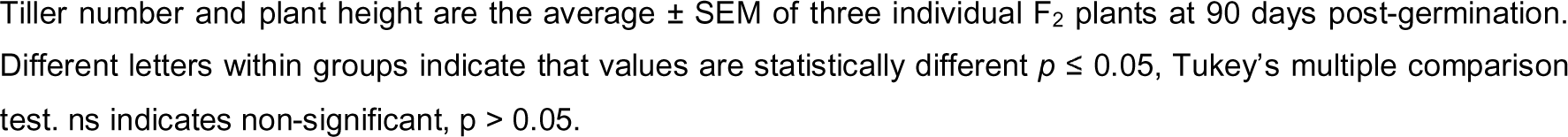
Leaf Chlorophyll content, plant height and tiller number of wild-type and OMT1/DiT2 double over-expressed lines.

|  | Chl (SPAD value) | Tiller number | Plant height (cm) |
| --- | --- | --- | --- |
| **Wild-type** | 43.8±1.1<sup>a</sup> | 17.3±1.6<sup>ns</sup> | 99.3±3.8<sup>ab</sup> |
| **OMT1-79/DiT2-44** | 39.9±1.6<sup>ab</sup> | 24.7±6.4<sup>ns</sup> | 97.7±1.8<sup>a</sup> |
| **OMT1-80/DiT2-39** | 40.9±1.6<sup>ab</sup> | 13±0.7<sup>ns</sup> | 96.7±3.1<sup>ab</sup> |
| **OMT1-45/DiT2-27** | 36.6±1.5<sup>b</sup> | 18±3.3<sup>ns</sup> | 87.7±4.7<sup>b</sup> |

Chl SPAD values are the average ± SEM of three leaves from three plants at mid-tillering stage using the upper fully expanded leaves.
Tiller number and plant height are the average ± SEM of three individual F<sub>2</sub> plants at 90 days post-germination. Different letters within groups indicate that values are statistically different $p \leq 0.05$ , Tukey's multiple comparison test. ns indicates non-significant, $p > 0.05$ .

**Fig. 10:**
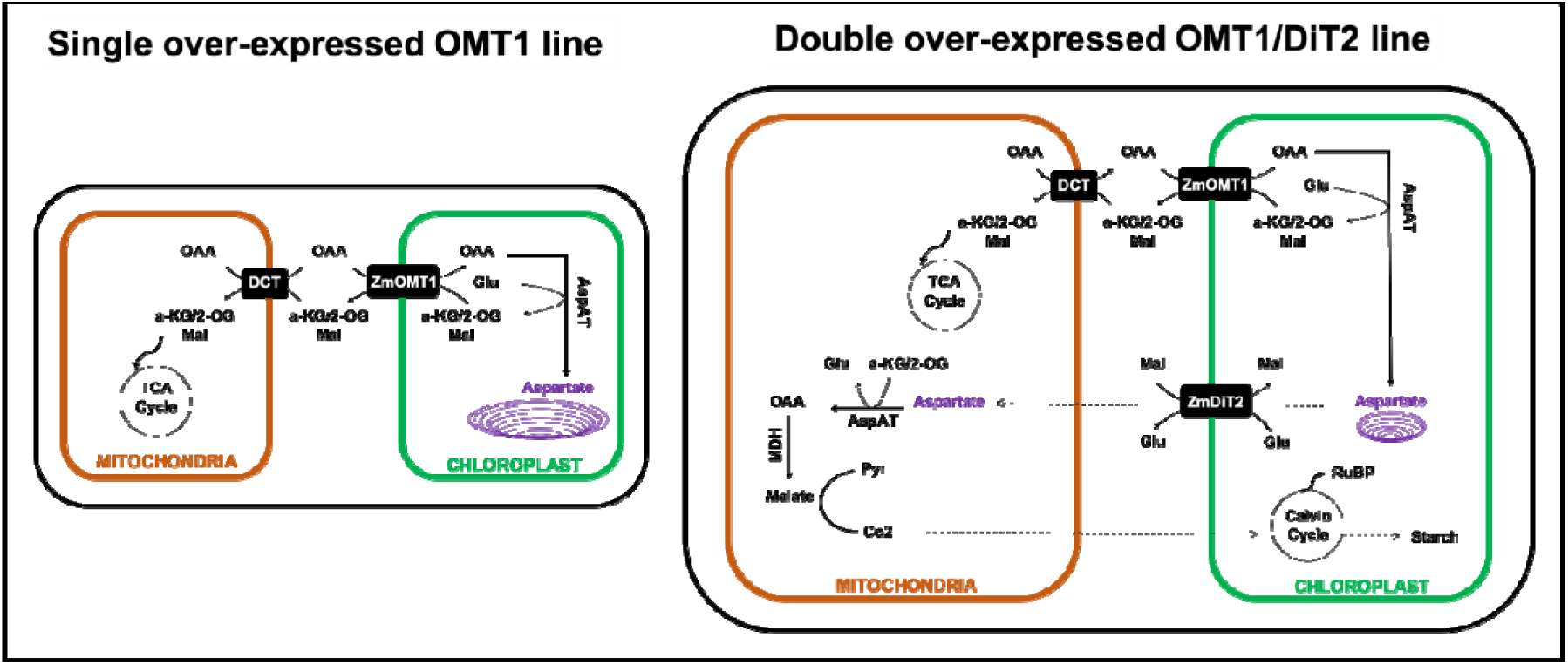
Schematic of the single OMT1 and double OMT1/DiT2 transgenic C_3_-rice plants. Both OMT1 and DiT2 were originated from C_4_-maize plant.

## Acknowledgements

This work was funded by the Bill and Melinda Gates Foundation C4-Rice project. Metabolite analyses were supported by the CEPLAS Plant Metabolism and Metabolomics laboratory, which is funded by the Deutsche Forschungsgemeinschaft (DFG, German Research Foundation) under Germany’s Excellence Strategy – EXC-2048/1 – project ID 390686111. We acknowledge the excellent technical assistance by E. Klemp, K. Weber, and M. Graf for GC-MS measurements. We would like to thank Prof. Jane Langdale (University of Oxford) for her C4-Rice team leadership and valuable comments on the manuscript draft. We wish to thank IRRI C4-Rice center for their help with plant transformation, husbandry and physiological measurements. We also wish to thank Sarah Covshoff for providing the constructs of OMT1 and DiT2.

